# Human NLRP1 and NLRP3 interact to drive inflammation in inflammosomopathies and inflammatory diseases

**DOI:** 10.64898/2026.01.26.701718

**Authors:** Juan Miguel Suarez-Rivero, Inés Muela-Zarzuela, Antonio Astorga-Gamaza, Rosa Antón, Sofía Bali, Joaquin Tamargo-Azpilicueta, Alejandra Guerra-Castellano, Raquel de la Varga-Martínez, Daniel García Cuesta, Irene Díaz-Moreno, Lukasz A. Joachimiak, Javier Oroz, Alberto Sanz, Mario D. Cordero

## Abstract

Inflammasomes are multiprotein complexes that form and activate after exposure to pathogenic microbes and host danger signals that trigger an inflammatory response. Although NLRP1 and NLRP3 inflammasomes share structural similarities and can be activated by similar stimuli, no evidence of heterotypic inflammasome assemblies has been reported. Here, we identify a unique interaction between NLRP1 and NLRP3 in human cells, forming a hybrid inflammasome, to drive inflammation. NLRP1 is essential for this hydrid inflammasome activation and NLRP3-mediated Caspase-1 activation and release of IL-1β and IL-18. The presence of the heterocomplex inflammasome was confirmed in blood samples from patients after kidney transplantation and is associated with inflammatory responses driven by NLRP3 and MEFV mutations that cause inflammasomopathies. Our findings reveal an unexpected level of intricacy in inflammasome composition, pinpointing hybrid targets that may pave the way for innovative pharmacological treatments for inflammatory disorders.

**Significance Statement:** Previous findings showed interactions between NLRP3 and NLRC4 or NLRP3 and NLRP11 showing that would be possible the interaction of the inflammasomes as supercomplexes and not working alone. Now, we show a new inflammasome heterocomplex inflammasome between NLRP1 and NLRP3 which is associated to the inflammatory profile in autoimmune diseases patietns and transplanted patients. These findings could open a new research topic for the design of dual inflammasomes inhibitors.

## Introduction

Inflammasomes are multiprotein complexes that assemble and activate in response to pathogenic microbes and host danger signals. Inflammasomes are indispensable for the auto-activation of Caspase-1, which mediates the maturation and release of IL-1β and IL-18. Caspase-1 activation can also induce an inflammatory form of cell death called pyroptosis. Pyroptosis dictates the organismal response to pathological processes, including chemotherapy and microbial infection (1). To initiate inflammasome complex formation, several cytosolic pattern recognition receptors (PRRs) act as sensors, detecting pathogen-associated molecular patterns (PAMPS) and damage-associated molecular patterns (DAMPS). The inflammasome family includes seven members of the NOD-like receptors (NLRs) (NLRP1, NLRP2, NLRP3, NLRP6, NLRP7, NLRP10, NLRP12 and NLRC4) and the HIN-200 family member, AIM2 (absent in melanoma 2), which uniquely recognizes and directly binds cytosolic DNA (2). Many inflammasomes utilize the adaptor protein apoptosis-associated speck-like protein containing a CARD (ASC) to facilitate the recruitment of Caspase-1 via CARD–CARD homotypic interactions.

NLRP3, which contains an amino-terminal pyrin domain (PYD), a central NACHT domain (domain present in NAIP, CIITA, HETE and TP1) and a carboxy-terminal leucine-rich repeat (LRR) domain, is one of the most well-studied inflammasomes in humans and mice. Genetic deletion of Nlrp3 in mice has been shown to extend lifespan and enhance health by delaying various age-related changes, including cardiac aging, insulin resistance, bone loss and ovarian aging (3–5). On the other hand, NLRP1 was the first NLR family member demonstrated to form an inflammasome complex, subsequently activating Caspase-1 (2). Still, despite its early discovery, NLRP1 remains one of the least understood members of the family. NLRP1 comprises an N-terminal PYD, a nucleotide-binding NOD/NACHT domain, five tandem LRR domains, a function-to-find domain (FIIND), and a C-terminal CARD.

NLRP1 and NLRP3 inflammasomes exhibit structural similarities and share common activators. Processes that trigger NLRP3 activation also engage in NLRP1, with both inflammasomes responding to equivalent PAMPs and DAMPs, such as ATP and K^+^ concentration. Besides, both NLRP1 and NLRP3 are also associated with a broad spectrum of human diseases including gain-of-function mutations (6,7). Moreover, a comprehensive review of the literature shows the participation of multiple inflammasomes, including NLRP1, NLRP3, NLRC4, NLRP6 or AIM2, in a variety of diseases, including neurodegenerative, cardiovascular, autoimmune diseases and cancer (Supplementary Table 1). This suggests the possibility of potential interactions or cooperative mechanisms among different inflammasomes. Previous reports indicated possible associations between NLRP3 and NLRP11 inflammasomes (8), between NLRP3 and NLRC4 (9) and more recently, between NLRP3 and NLRP12 (10). However, a direct binding between NLRP1 and NLRP3 inflammasomes has never been shown. Considering the significance of NLRP1 and NLRP3 in human diseases, a direct interaction between them would reveal a more nuanced and multifaceted mechanism underlying inflammasome assembly and regulation. Heterocomplexes formed by combining two distinct inflammasomes would integrate the properties of each individual complex while introducing novel features and functions similar to mitochondrial respiratory supercomplexes (11).

To the best of our knowledge, direct contact between NLRP1 and NLRP3 has not been described before. We propose that a new framework—considering the existence of inflammasome heterocomplexes—offers two significant advantages: a deeper understanding of the role inflammasomes play in human diseases, and the potential development of dual inhibitors that could enhance therapeutic efficacy.

This study demonstrates the unprecedented formation of hybrid NLRP1 and NLRP3 inflammasomes in human cells. We show that the N-terminal domains of NLRP1 can bind to NLRP3’s NACHT domains using mass spectrometry. These contacts are instrumental in activating the inflammasome heterocomplex and cytokine release. In the absence of NLRP1, NLRP3-mediated Caspase-1 activation and the release of IL-1β and IL-18 are impaired. Conversely, inhibition of NLRP3 triggers a compensatory overactivation of NLRP1, leading to effective Caspase-1 activation and cytokine release. We confirmed the physiological significance of this hybrid inflammasome formation in human samples from kidney transplant patients. Notably, these samples exhibited elevated cytokine release that persisted despite immunosuppressive treatment, likely because the inflammasomes heterocomplex were not effectively targeted. Strikingly, both inflammasome heterocomplex formation and LPS-induced inflammation were diminished in CAPS (Cryopyrin-Associated Periodic Syndromes) and Familial Mediterranean Fever (FMF) patients following NLRP1 silencing. Overall, our findings establish novel paradigms in inflammasome activation and pave the way for innovative pharmacological strategies to treat inflammatory diseases.

## Results

### NLRP1 interacts with NLRP3 in human cells

To assess the role of NLRP1 in NLRP3 inflammasome activation, we used skin fibroblasts from a patient harbouring a pathological NLRP3(p.Q703K) gain of function mutation (12). These fibroblasts showed co-expression of both NLRP1 and NLRP3 proteins (Figure 1A). Proximity Ligation Assay (PLA) experiments demonstrated the proximity of NLRP1 with NLRP3 in mutant fibroblasts at basal levels (Figure 1B). Subsequently, we extended our study to investigate this novel interaction in an immune cell type, comparing PLA data in THP1-derived macrophages exposed to the classical inflammasome inducer lipopolysaccharide (LPS)+ATP and Nigericin. NLRP1 and NLRP3 exhibited proximity in macrophages following LPS+ATP and nigericin stimulation, as indicated by numerous red dots (positive PLA) confined within the cellular boundaries (Figure 1C). To mitigate potential false-positive signals, we stimulated differentiated macrophages derived from THP-1 wild-type, NLRP1 knockout, and NLRP3 knockout cells and then stained them with anti-NLRP1 and anti-NLRP3 antibodies for PLA experiments. Our findings revealed that the presence of both proteins was required to detect interactions. Similarly, double fluorescence imaging demonstrated positive colocalization between NLRP1 and NLRP3 and the independent localization of each protein (Figure 1D).

**Figure 1.**
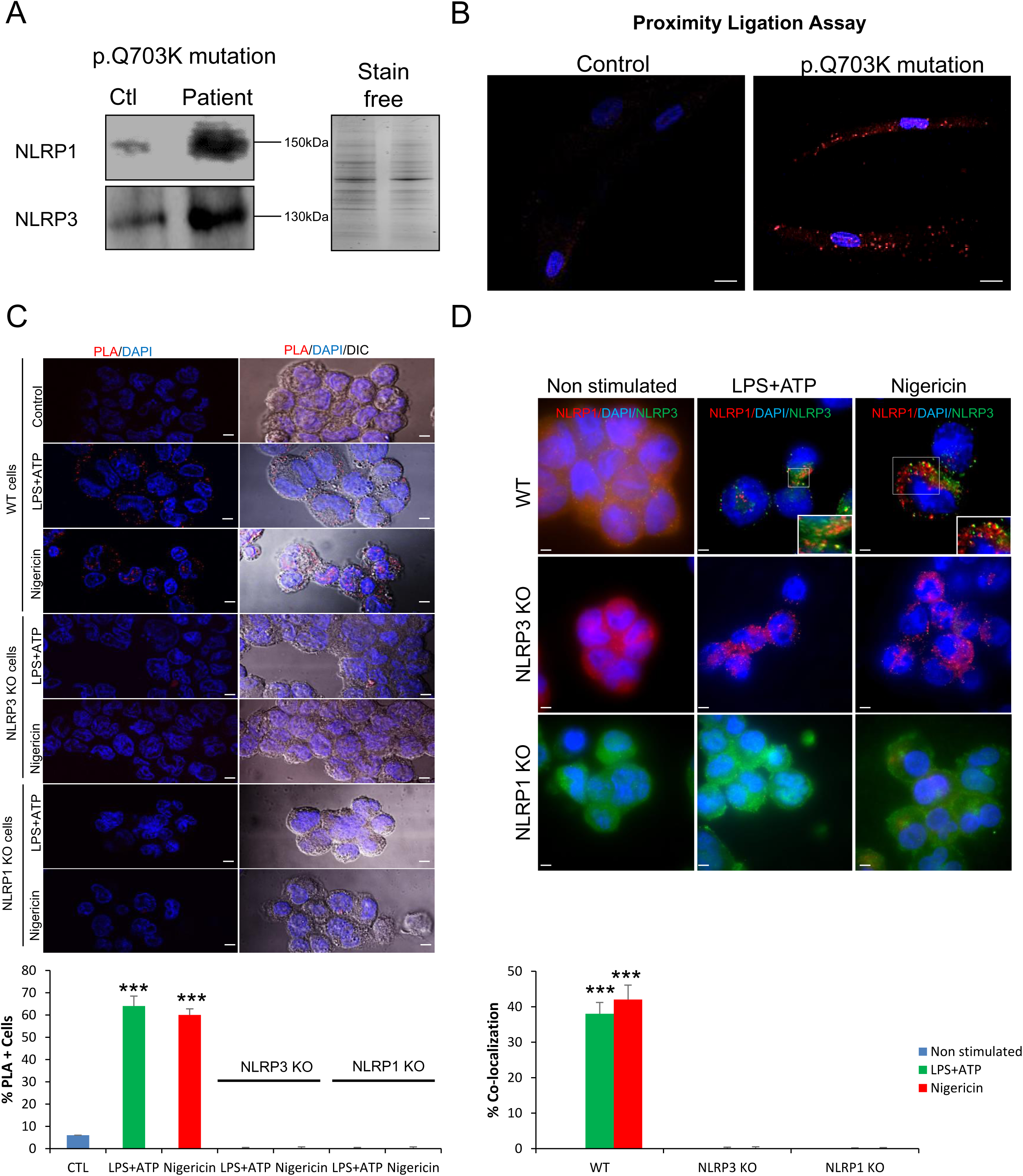
NLRP1 interact with NLRP3 in human cells. **A.** NLRP1 and NLRP3 protein expression in skin fibroblasts from a patient with a p.Q703K mutation compared with healthy fibroblasts (Ctl) as monitored by immunoreactivity with NLRP1 and NLRP3 antibodies. Stain free SDS-PAGE gel is shown on the right. **B.** Representative microscopy images of fibroblasts from patient and control individuals stained with anti-NLRP1 and anti-NLRP3 antibodies for PLA experiments. Nuclear stain is obtained with DAPI. The scale bars represent 10 μm. **C.** Representative microscopy images of macrophages stained with anti-NLRP1 and anti-NLRP3 antibodies for PLA experiments stimulated with LPS+ATP or Nigericin. Nuclear stain was obtained with DAPI. The scale bars represent 10 μm. **D.** Representative microscopy colocalization images of NLRP1 and NLRP3 antibodies in macrophages stimulated with LPS+ATP or Nigericin. Nuclear stain is obtained with DAPI. The scale bars represent 10 μm. NLRP1 or NLRP3 KO cells were used as negative controls. Data is presented as the mean ± SD of three independent experiments. (***p < 0.001).

**Figure 2.**
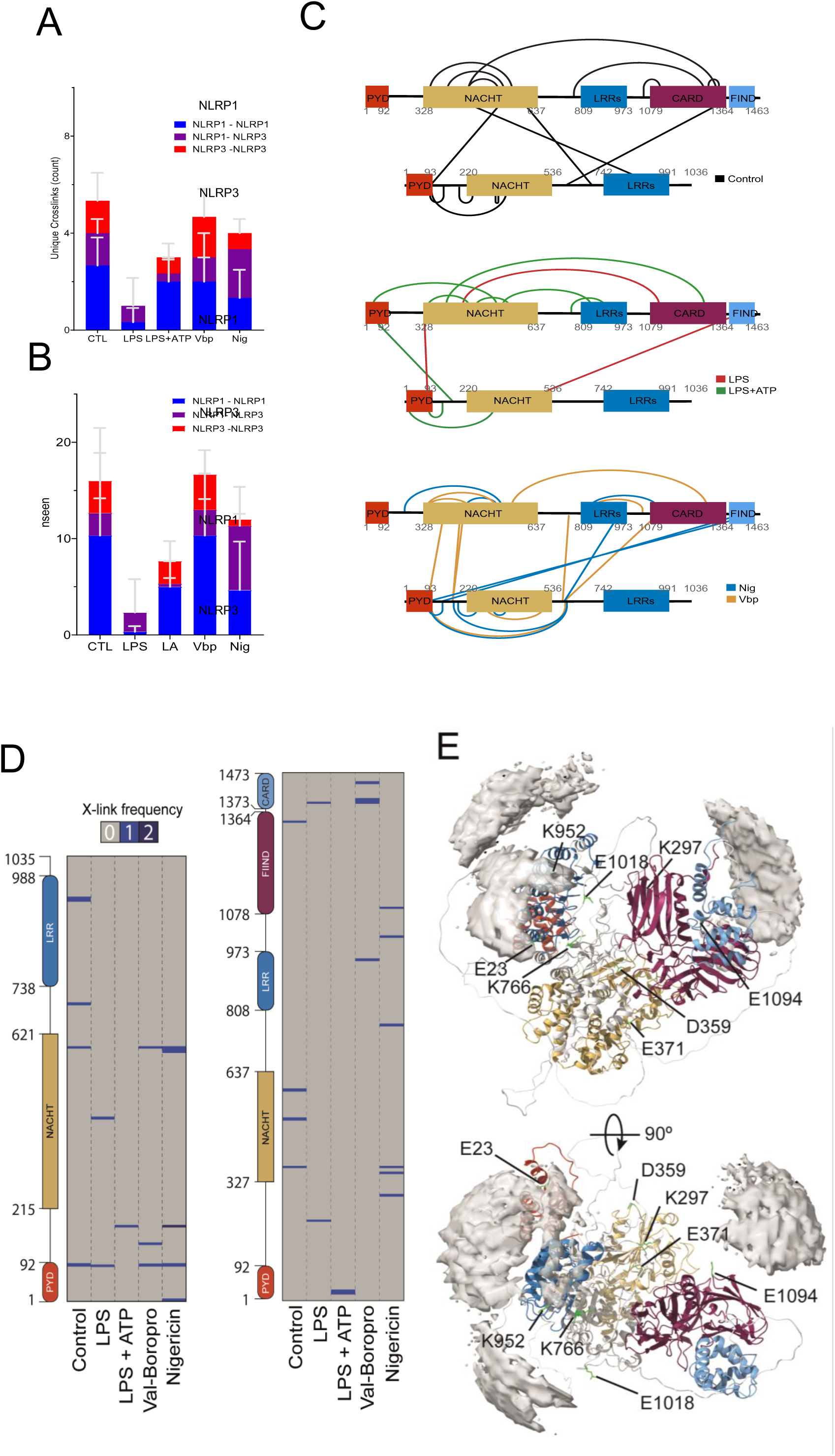
Topology of the inflammasome heterocomplex. **A.** Unique contacts between NLRP1-NLRP1 (blue), NLRP1-NLRP3 (purple) and NLRP3-NLRP3 (red) observed across three biological replicates. **B.** Average abundance of contacts between NLRP1-NLRP1 (blue), NLRP1-NLRP3 (purple) and NLRP3-NLRP3 (red) observed across three biological replicates for Control (CTL), Lipopolysaccharide (LPS), Lipopolysaccharide+ATP (LPS+ATP), ValboroPro (VbP) and Nigericin (Nig) stimulated samples. Data is reported as averages with standard deviation (n=3). **C.** Contact maps illustrating the domain distribution of unique cross-links betweeen NLRP1-NLRP1, NLRP3-NLRP3 and NLPR1-NLRP3 for CTL (black), LPS (red), LPS+ATP (green), vbp (yellow) and Nig (blue). The PYD, NACHT, LRRs, CARD and FIND domains are colored red, yellow, blue, purple and light blue. **D.** Cross-linking-derived NLRP3-NLRP1 identified peptides across NLRP3 (left) and NLRP1 (right) sequence under the different treatments. **E.** The full-length structure of NLRP1 deposited in Alphafold structure prediction database (domains colored according to the color code of panels C and D) and the reduced accessible interaction space of NLRP3 consistent with at least two cross-links (grey surface), as predicted by DisVis Webserver using the residues identified in the crosslinking of cells treated with the inflammasome inductors (LPS+ATP, Val-boroPro (VbP) and Nigericin).

Immunoprecipitations of both NLRP1 and NLRP3 further confirmed the interaction between NLRP1 and NLRP3 in macrophages treated with Nigericin (Supplementary Figure 1A). As excpected, NLRP1 and NLRP3 KO cells showed negative results. Native PAGE also showed the formation of an NLRP3-dependent high molecular mass complex in WT macrophages stimulated with LPS+ATP (LA) or Nigericin (Nig) (Supplementary Figure 1B) (Supplementary figure 2; specificity of the antibodies was showed using HEK293T cells with mutant NLRP3). These high molecular weight complexes, which were absent in NLRP1 or NLRP3 KO cells, contained both NLRP1 and NLRP3, in agreement with the formation of a hybrid inflammasome. Since proteins remain under non-denaturing conditions, the band patterns provide information on both free and associated complexes (Supplementary Figure 1B). The protein composition of the upper and lower bands showed from the Native PAGE was analyzed using mass spectrometry and identified through tryptic digestion. Overall, the results provide evidence that NLRP1 and NLRP3 interact to form a hybrid inflammasome (Supplementary Figure 3A and B and Supplementary Table 2).

### Topology of the inflammasome heterocomplex

To understand the molecular interactions of NLRP1 under various stimuli (LPS+ATP, VbP and nigericin which can induce both NLRP1 and NLRP3 activation), we performed NLRP1 IP using specific antibodies and crosslinked the reaction using a zero-length crosslinker, 4-(4,6-Dimethoxy-1,3,5-triazin-2-yl)-4-methylmorpholinium chloride (DMTMM). This crosslinker activates acidic residues predominantly Asp and Glu acids - enhancing their reactivity towards the primary amines found mainly on lysines, thereby exposing charge-complementary electrostatic surfaces. The IP’s were reacted with DMTMM, quenched and processed using our crosslinking-coupled to mass spectrometry (XL-MS) pipeline (13). To quantify the reproducibility of the observed contacts, we interpreted the observed crosslinks as a function of three biological replicates. Specific crosslinks between NLRP1-NLRP1, NLRP3-NLRP3 as well as heterotypic contacts between NLRP1-NLRP3 were identified. We first interpreted the contacts as the average number of observable unique contacts between the different pairs across the replicates (Figure 4A and B). We found that in the LPS condition, fewer sites between all pairs of proteins were detected with a notable absence of NLRP3-NLRP3 contacts. Stimulation with LPS and ATP recovers NLRP3-NLRP3 contacts towards numbers of observed contacts in Vbp and nigericin. Mapping of the detected intra- and intermolecular, unique contacts on NLRP1 and NLRP3 domains highlight contacts between NLRP1 NACHT and NLRP3 C-term in the control condition (Figure 4C, black), while treatment with nigericin and Vbp change the overall arrangement of contacts between NLRP1 and NLRP3 (Figure 4C, blue and yellow), underscoring the inherent flexibility of NLRP1 and NLRP3 domains. As observed in the overall quantification, the LPS condition (Figure 4C, red) dramatically loses NLRP1 and NLRP1-NLRP3 contacts and leads to a complete loss of NLRP3 contacts. Finally, the addition of LPS+ATP shows a recovery of these contacts (Figure 4C, green). This data suggests that addition of LPS reduces NLRP1 assembly and recruitment of NLRP3 and the addition of LPS+ATP recovers NLRP1 assembly and recruitment of NLRP3.

The NLRP1 N-terminal proteasomal target region was found to interact with NLRP3 primarily through NLRP3’s PYD and NACHT domains upon induction with inflammasome inductors (Figure 4D). To obtain further insights into the NLRP1-NLRP3 complex formation, residues identified in XL-MS experiments on the NLRP1 N-terminal PYD-NACHT-LRR-ZU5 domains (residues 1-1213) and their corresponding counterparts in full-length NLRP3 were used as intersubunit distance restraints to define the accessible interaction space (Figure 4E). A comprehensive exploration of 8,060,776,715 complexes yielded 99,387 possible orientations according to at least two simultaneous XL-MS restraints (out of 8). The centers of mass of NLRP3 were grouped in three defined patches: two of them majorly encompassing the back side of the LRR domain, and one of them closer to the FIIND domain. Overall, structural and biophysical data indicate that an hybrid inflammasome between NLRP1 and NLRP3 is associated by different interaction between NACHT and LRR domains.

### Temporal dynamics of hybrid inflammasome assembly in immune cells

Different stimuli have been used to activate inflammasome complexes *in vitro* with varying time dependencies. For example, cells are treated with LPS for 4 h and subsequently with ATP for 30 min to induce NLRP3-inflammasome activation (14), and nigericin, which is known to stimulate both NLRP3 and NLRP1 (20), shows activation as early as 1 h after application (15, 16). Given these variations, we investigated whether hybrid inflammasome assembly exhibited similar time progression. THP1-derived macrophages were stimulated with LPS and ATP (following a proportional time of 88% of the time LPS and 12% of the time ATP) and the activation of NLRP1 and NLRP3 was monitored over 1, 2, 3 and 4 h via SDS and Native PAGE. Parallel experiments were conducted with nigericin where NLRP3 activation was observed from the beginning, with NLRP1 incorporation noted after 45 min (Figure 3A and Supplementary Figure 4). However, both proteins levels were significantly reduced by 120 min of nigericin treatment, suggesting potential complex disassembly (Figure 3A and Supplementary Figure 4). Remarkably, there is a significant correlation of IL-1β cytokine secretion in macrophages induced by LPS+ATP and nigericin with hybrid supercomplex formation (Figure 3B), indicating a clear pro-inflammatory role for the NLRP1:NLRP3 inflammasome heterocomplex.

**Figure 3.**
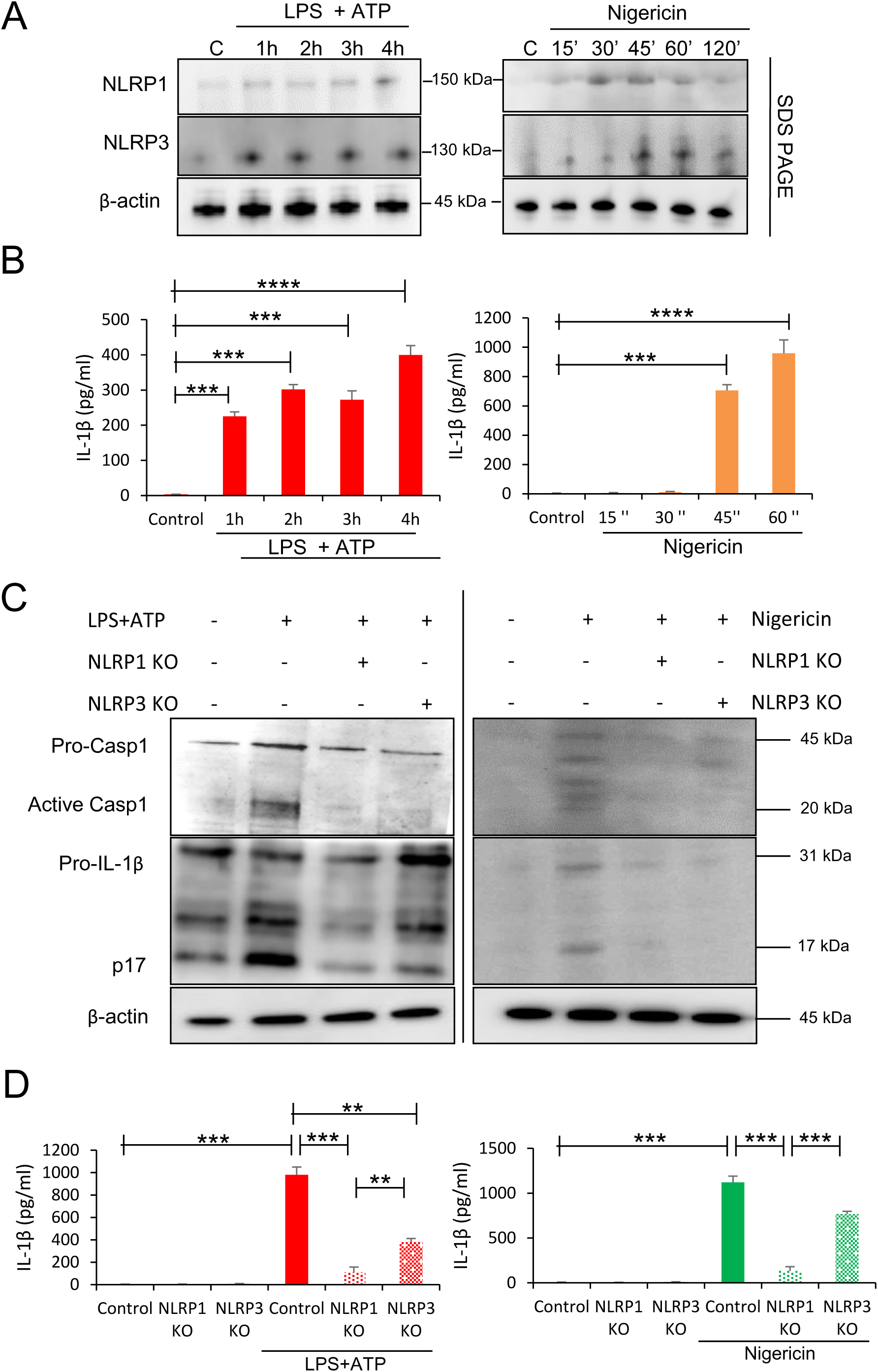
Hybrid inflammasome assembly is time-dependent. **A.** Western blots on SDS of macrophages treated with three different inflammasome inductors (LPS+ATP and Nigericin) and isolated at different times. Antibodies used for blotting are specified on the left. **B.** IL-1β release after the different time of treatments (describer, top, bottom panel) in culture supernatants of macrophages as monitored by ELISA. **C.** Immunoblots of cell lysates from NLRP1 and NLRP3 KO macrophages treated with LPS+ATP or Nigericin. Antibodies used in immunoblots are specified. **D.** IL-1β release in culture supernatants of macrophages as monitored by ELISA. Data is presented as the mean ± SD of three independent experiments. (**p < 0.01; ***p < 0.001; ****p < 0.0001).

### NLRP1 is a key regulator of the inflammasome heterocomplex

To evaluate the functional impact of each inflammasome in the activation of the heterocomplex, we stably KO NLRP1 cells by using Crispr/Cas 9 in THP1 cells followed by differentiation of the cells into macrophages. Furthermore, NLRP3 KO THP1 cells were kindly gifted by Dr. Pablo Pelegrin (IMIB-Arrixaca, Murcia). We utilized various stimuli (LPS+ATP and nigericin) to induce inflammasome activation and measured the levels of NLRP1, NLRP3, Caspase-1 and IL-1β. NLRP1 or NLRP3 KO reduced the levels of activated hybrid inflammasome formation and Caspase-1 and IL-1β release (Figure 3C and D). Notably, the inhibition of NLRP1 was more effective reducing the activation of the hybrid inflammasome and demonstrated a more significant inhibition of the IL-1β secretion (Figure 3D). However, NLRP1 KO also induced a reduction in NLRP3 protein expression and activation. To ensure sgRNA specificity of NLRP1, a nucleotide Basic Local Alignment Search Tool (BLAST) analysis was performed on the datasheet’s target sequences (Supplementary Table 3) showing specific effect in NLRP1 and not in NLRP3. Therefore, this inhibitory effect could be showing unknown possible new mechanisms of interaction between the different inflammasomes.

Finally, these findings underscore the critical role of NLRP1 in regulating NLRP3 and the hydrid inflammasome, highlighting its potential importance in the design of pharmacological strategies.

### The role of the inflammasome heterocomplex in human inflammatory diseases

Inflammasome complexes have been implicated in inflammatory processes across a range of human diseases, including cardiovascular, autoimmune, metabolic, and neurodegenerative disorders, thereby contributing directly or indirectly to their pathophysiology (3–7). Given the known role of inflammasomes in disease progression, hybrid inflammasomes are similarly expected to contribute to and potentially exacerbate inflammatory processes within disease contexts.

One example of such increased and prolonged inflammation are the inflammasomopathies. To investigate the role of the NLRP1-NLRP3 inflammasome heterocomplex in the context of these genetic diseases, we assessed blood samples from a patient with CAPS who carries the germline pathogenic NLRP3 gain-of-function variant p.S726G (Supplementary Table 4). Serum levels of IL-1β were similar to those of healthy controls, albeit with moderately elevated IL-18 levels (Figure 4A). Cultured PBMCs from these patients, both with and without LPS stimulation, showed increased levels of IL-1β and IL-18, as well as LDH, compared to healthy individuals (Figure 4B and C). Western blotting analysis showed increased expression of NLRP1, NLRP3, Caspase-1 activation, and cleaved gasdermin D (GSDMD) following LPS stimulation (Figure 4D). Interstingly, native PAGE experiments also revealed the presence of the hybrid inflammasome in the circulating blood of these patients (Figure 4E). NLRP1 knockdown reduced the inflammasome heterocomplex assembly, Caspase-1 activation, and reduced IL-1β, IL-18 and LDH levels (Figure 4A-E). The inflammasome heterocomplex was also detected in PBMCs from four patients with FMF, who carried mutations in the MEFV gene (p.G632S, 1 patient; p.M694V, 1 patient; and the duplication c.383_391dup, p.E128_N130 dup, 2 patients) (Supplementary Table 5), with increased formation after LPS treatment (Supplementary Figure 5A). Remarkably, this hybrid inflammasome increase was also associated with an elevated release of IL-1β and IL-18 (Supplementary Figure 5B). These findings suggest a significant role for the NLRP1:NLRP3 inflammasome heterocomplex in inflammasomopathies, promoting inflammation in response to different stressors.

**Figure 4.**
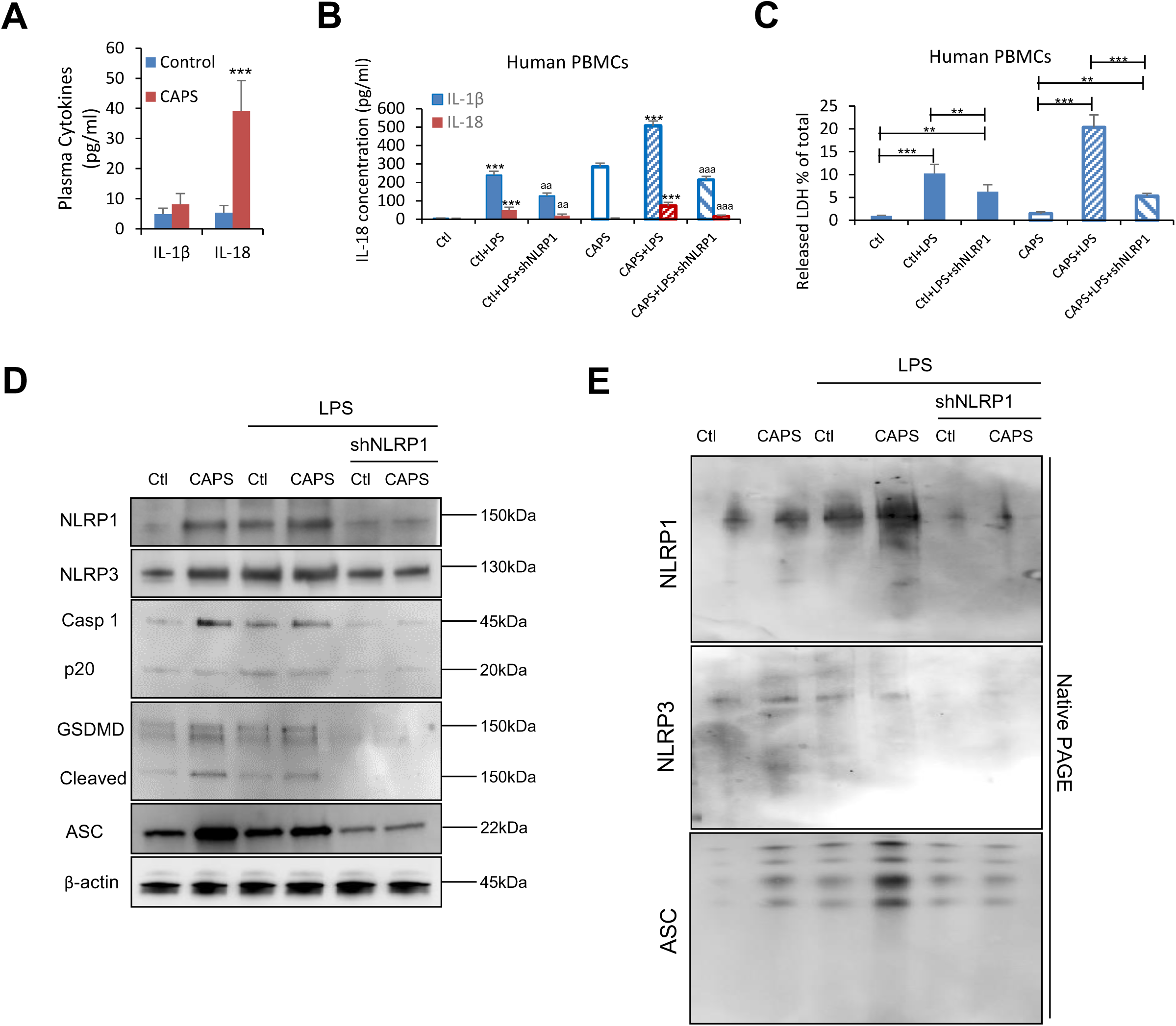
Hybrid inflammasome are formed in human diseases. **A.** IL-1β and IL-18 serum levels from patients with NLRP3 mutations. **B and C.** IL-1β, IL-18 (B) and LDH (C) release from PBMCs after treatment for 4h with LPS (100 ng/ml) and treated with NLRP1 shRNA (shNLRP1) (n = 5 HD, n = 3 CAPS patients) as monitored by ELISA. Data is presented as the mean ± SD of three independent experiments (***p < 0.001, **p < 0.01 LPS *vs* no LPS; ^aaa^p < 0.001, ^aa^p < 0.01 LPS *vs* LPS+shNLRP1). **D and E.** Immonublot analysis on SDS (**D**) and native (**E**) PAGE using the specified antibodies (left) on lysated PBMCs from a representative healthy (Ctl) and CAPS patient.

Finally, other example of such increased and prolonged inflammation is observed in organ transplantation scenarios. We analyzed samples from kidney-injured patients before and after transplantation to investigate whether the NLRP1-NLRP3 inflammasome heterocomplex is active in such a pathological context (Supplementary Table 5). Peripheral blood mononuclear cells (PBMCs) were isolated 24 h prior to, 72 h, and 7 days after the transplant. Native PAGE experiments conducted on these samples indicated the presence of the hybrid inflammasome in the circulating blood before and after the renal transplant (Supplementary Figure 6A-D). Interestingly, one-week post-transplant, the patients displayed elevated levels of the NLRP1-NLRP3 inflammasome heterocomplex. Flow cytometry analysis identified a monocyte population characterized by ASC specking and an intense co-expression of NLRP1 and NLRP3 (Supplementary Figure 6E). Notably, this population was significantly enriched between 72 h and seven days post-transplant compared to the pre-transplant levels (Supplementary Figure 6F). Accordingly, we observed increased IL-1β serum levels in these patients (Supplementary Figure 6G). Importantly, all patients received standard immunosupressant treatments (tacrolimus, mycophenolate mofetil, steroids and basiliximab/thymoglobulin) before to transplantation to minimize immune responses and enhance transplant success. Despite of the demonstrated inhibitory effects of tacrolimus on NLRP3 in experimental models (16, 17), our findings support activated inflammatory response for these patients via inflammasome heterocomplex formation.

## Discussion

NLRP1 was the first NLR to be identified; however, it remains one of the least well-understood members of the family (18). Although NLRP1 has been implicated in numerous human diseases, the precise mechanisms remain unclear. For many years, its activation was thought to be limited to pathogen encounters and infection-associated immune responses. However, physiological activators of human NLRP1 have not been extensively characterized (19). Recent studies have begun to identify the danger activation signals of NLRP1 and shed light on its functional relevance. Here, we demonstrated that NLRP1 interacts with NLRP3 to form a superstructure, which could be complatible with a supercomplex. This supercomplex is crucial for activating, sustaining, and amplifying the inflammatory response in human macrophages. NLRP1 interaction with NLRP3 was required for Caspase-1 activation, release of IL-1β and IL-18 and pyroptosis.

Inflammasome complexes have traditionally been studied as isolated units responsible for inflammatory responses. However, structural and functional similarities enable homotypic protein interactions among these multiprotein complexes. Previous interactions have been reported, including NLRC4-NLRP3 (20,21), NAIP-NLRC4 (22,23), and NLRC5 modulating NLRP3 through an unknown mechanism (24). These studies, however, have only yielded initial observations. Notably, previous research demonstrated complex formation between Nod2 and NLRP1, where inhibiting NLRP1 also reduced IL-1β activation, supporting our findings with the NLRP1-NLPR3 supercomplex (25).

In this study, we outline the timeline of supercomplex assembly to demonstrate the interaction between NLRP1 and NLRP3. Our results illustrate the dynamic process of their association and dissociation, which corresponds with cytokine release. Additionally, we described the pathophysiological roles of these hybrid inflammasomes in two distinct inflammatory processes, kidney transplanted patients and individuals with inflammasome gain-of-function mutations, which could be extrapolated to other human inflammatory diseases.

Acute inflammation refers to how innate immunity fights infections and helps speed up the healing process, which would be associated with our *in vitro* experiments with typical inflammasome inductors (LPS+ATP, VbP, Nigericin) (19). In this scenario, the hydrid inflammasome may serve as a platform that amplifies inflammatory responses, with NLRP1 playing a critical role—as evidenced by the marked inhibition observed following its silencing.

This concept of an inflammation-enhancing platform is plausible in human diseases where infections are accompanied by cell damage and metabolic dysregulation. In line with our findings, Oh and colleagues recently proposed that simultaneous activation of multiple inflammasomes by diverse ligands can trigger various downstream effects, including inflammatory cell death (26). They exposed macrophages to LPS+ATP, Flagellin, poly(dA:dT) and TcdB, which activated at the same time the NLRP3, AIM2, NLRC4, and Pyrin inflammasomes (27). Since NLRP1 and NLRP3 are known to detect various signals associated with cell damage and infection, we demonstrate a direct molecular interaction between these two inflammasome sensors. This interaction drives the assembly of an inflammasome heterocomplex that amplifies inflammation and promotes cell death. Moreover, in the context of chronic inflammation, which is linked to long-term human diseases, an hybrid inflammasome could also explain the cycle of inflammation, stabilization, and aggravation of the symptoms which are typically observed in these conditions.

Finally, our findings and previous observations of interactions between different inflammasomes suggest the need to reconsider therapeutic strategies. Merely inhibiting a single inflammasome may result in the compensatory activation of others. Therefore, a more comprehensive approach that addresses the complex network of inflammasome is required.

## Material and Methods

### Biological samples

Study protocols were approved by the corresponding Ethical Committee of the Hospital Puerta del Mar, Cádiz Spain. Peripheral blood mononuclear cells (PBMCs) we obtained from blood samples from patients with inflammasomopathies (Supplementary Table 4) and chronic kidney disease (Supplementary Table 5) and, before and after being subjected to renal transplantation. All subjects included in the study provided written informed consent.

Peripheral blood monocytes (PBMC) and serum were obtained from patient blood using Cytiva Ficoll-Paque™ PLUS (Thermo Fisher, Waltham, MA, USA, 11768538). Commercial THP-1 cell line was purchased from ATCC (Manassas, VI, USA, TIB202).

### Cell culture

PMBC’s and THP-1 cells were seeded in 6 wells plates in RPMI medium supplemented with 10% FBS and 1% antibiotics (Thermo Fisher, Waltham, MA, USA, 11548876), and kept at 37°C in a CO_2_ 5% incubator.

THP1 cells were differentiated into macrophages using phorbol-12-myristate-13-acetate (PMA) at 50 nM for 24 hours. Macrophages were scrapped for protein extraction, and medium aliquoted for ELISA and LDH analysis.

### *In situ* Protein Ligation Assay (PLA)

Skin fibroblasts or THP-1 derived macrophages were grown on 1 mm width glass coverslips for 72 hours in high glucose DMEM medium containing 10% FBS and 1% antibiotics. They were washed twice with PBS, fixed in 3.8% paraformaldehyde for 15 min at room temperatura (RT), permeabilized with 0.1% Triton X-100 in PBS for 10 min and incubated in blocking buffer (BSA 1%, Triton X-100 0.05% in PBS) for 30 min. In the meantime, the primary antibody was diluted 1:100 in antibody buffer (BSA 0.5%, Triton X-100 0.05% in PBS). Macrophages were incubated 2 h at 37°C with the primary antibody and subsequently washed twice with PBS. The secondary antibody was similarly diluted 1:400 in antibody buffer and their incubation time with cells was 2 h at RT. PLA experiments utilized DuoLink (Sigma-Aldrich DUO92007): after washing primary antibodies, cells were then incubated with the appropriate probes (Sigma Aldrich DUO92004 and DUO92002) for 1 h at 37 °C and washed two times. Probes were then ligated for 30 min at 37 °C, washed two times in Buffer A and amplified using the manufacturer’s polymerase for 100 min at 37 °C in the dark. Coverslips were washed twice with PBS, incubated for 5’ with PBS containing DAPI (1 µg/ml) and washed again with PBS. Next, they were mounted on microscope slides using Vectashield Mounting Medium (Vector Laboratories, Burlingame, CA, USA, H1000).

### Coimmunoprecipitation

THP1 cells were seeded in T75 flasks until 90% confluency. Each cell pellet was resuspended in native cell lysis buffer. After 30 min of incubation in ice, cell lysate was centrifuged 2’, 12000g, 4 °C. Protein concentration was measured using Lowry Protein Assay (Biorad, Hercules, CA, USA, 5000112). Antibodies were added in the ratio stated in the manufacturers data sheet. Protein-antibody mixture was incubated overnight at 4°C in subtle rotation. For protein purification, protein A magnetics beads were used (Thermo Fisher, Waltham, MA, USA, 88845). Magnetics beads were washed twice with PBS-Tween (0,005%) and incubated alongside the mixture protein-antibody for 1 h at 4 °C under rotation. Beads were washed three times with PBS-Tween and once with deionized water. Input was stored as immunoprecipitation control. For the elution step, 100 μL of soft elution buffer (0,2% SDS, 0,1 % Tween 20 dissolved in 50 mM Tris-HCl pH 8) were added to the beads and incubated during 10 min at RT in agitation. Samples were magnetized and supernatant was transferred to a new Eppendorf for further analysis.

### Protein electrophoresis

For SDS-PAGE, proteins were extracted using RIPA buffer (Thermo Fisher, Waltham, MA, USA, 89901), preserved in LDS NuPAGE buffer (Thermo Fisher, Waltham, MA, USA, NP0008) and heatshocked at 95°C for 2 min. Gel electrophoresis was performed using 4–20% Mini-PROTEAN® TGX Stain-Free™ Protein Gels (Biorad, Hercules, CA, USA) at 200V in Tris-Glycine-SDS buffer (Biorad, Hercules, CA, USA, 1610772) for 45 min.

For native PAGE, proteins were extracted using native cell lysis buffer (Abcam, Cambridge, UK, ab156035) and preserved in Tris-Glycine sample buffer (Thermo Fisher, Waltham, MA, USA, LC267). Gel electrophoresis were performed using 4–20% Mini-PROTEAN® TGX Stain-Free™ Protein Gels (Biorad, Hercules, CA, USA) at 200V in Novex Tris-Glycine buffer (Thermo Fisher, Waltham, MA, USA, LC2672) for 4 hours in ice.

### Western Blotting

Protein transfer was made using a TurboTransfer (Biorad, Hercules, CA, USA) at 25V for 10 min. After transferring the proteins to 0.45 µM nitrocellulose membranes (Biorad, Hercules, CA, USA), these were incubated for 1 h in BSA 5% in PBS-Tween20 0.05% and then overnight at 4°C with primary antibodies at 1:1000. Then washed twice with PBS-Tween20 and incubated with the corresponding secondary 1:5000 antibody for 1 h at RT. Protein loading was checked using stain free gel activation and reference genes. Stripping was not used.

### Flow cytometry

PBMCs were stained with LIVE/DEAD Fixable Aqua Dead Cell Stain (Thermo Fisher, Waltham, MA, USA, L34966) for 20 min at RT. After washing once with PBS, cells were fixed and permeabilized with Fixation/Permeabilization Solution (Becton Dickinson, Franklin Lakes, NJ, USA, 554714) for 20 min at 4°C and washed with BD Perm/Wash buffer. Then, cells were stained with antibodies for 20 min at RT. Finally, cells were washed with BD Perm/Wash buffer. Samples were acquired on a BD FACSCelesta flow cytometer and data was analyzed using FlowJo V10 software. Gating was performed according to the different fluorescence minus one (FMO) control.

### Interleukin measurement

ELISA kits IL-18 (ab215539), and IL-1β (ab214025) were purchased from Abcam (Cambridge, UK). Assays were performed using patient blood serum or cell medium according manufacturers protocol.

### LDH assay

LDH/ Lactate Dehydrogenase Assay Kit (Colorimetric) (Abcam, Cambridge, UK, ab102526) was performed using 50µL of cell medium according manufacturers protocol.

### CRISPR/ Cas9

NLRP1 was knocked out in cells using the NLRP1 sgRNA CRISPR/Cas9 All-in-one Lentivector set for humans (Applied Biological Materials, Richmond, Canada, 3190611). To ensure sgRNA specificity, a nucleotide Basic Local Alignment Search Tool (BLAST) analysis was performed on the datasheet’s target sequences (Supplementary Table 3).

Cells were transduced via spinoculation as follows: a 1 mL suspension containing 10^6^ THP1 cells was mixed with 1 μl/ml of each lentivirus stock and 8 μg/mL polybrene (Santa Cruz Biotechnology, Santa Cruz, CA, USA, sc-134220), then incubated at room temperature for 20 min. The mixture was centrifuged at 800 × g at 37°C for 30 min. After centrifugation, the supernatant was discarded, and the cells were seeded in 6-well plates. After 72 h, cells were selected using 1 μg/mL puromycin (Santa Cruz Biotechnology, Santa Cruz, CA, USA, sc-108071). Knockout-positive cells were confirmed by qPCR and western blotting

### Reagents

Bovine Serum Albumim (A7030), Triton X-100 (X100-100ML) and PMA (P8139) were purchased from MERCK (Darmstadt, Germany). The following reagents were obtained from Thermo Fisher Scientific (Waltham, MA, USA): RPMI 1640 medium (31870074), NativeMark™ Unstained Protein Standard (LC0725), silencer select siRNA NLRP1 (4392420 id: s22521) and silencer select siRNA NLRP3 (4392420 id: s41556). Primary antibodies: anti ASC antibody (Abcam, Cambridge, UK, ab283684), anti-Caspase 1 antibody (Cell Signaling, Danvers, MA, USA, 3866), anti C-terminal NLRP1 antibodies (Abclonal, Woburn, MA, USA, A16212; Cell Signalling Danvers, MA, USA, 56719S), anti NLRP3 antibody (Abclonal, Woburn, MA, USA, A5652), anti β actin antibody (Abcam, Cambridge, UK, ab6276), anti N-terminal NLRP1 antibody (Novus Bio, CO, USA, NBP1-54899), anti GAPDH antibody (Cell Signaling, Danvers, MA, USA, 5174S), anti interleukin 1β antibody (Cell Signaling, Danvers, MA, USA, 12703) and, anti gasdermin antibody (Abcam, Cambridge, UK, ab219800)

Flow cytometry antibodies: The following antibodies were obtained from Becton Dickinson (Franklin Lakes, NJ, USA) anti-CD56-FITC (B159, 562794), anti-CD3-PE-Cy7 (SK7, 557851), anti-CD14-APC-H7 (M5E2, 561384), and anti-CD4-BV605 (RPA-T4, 562658). Anti-HLA-DR-PE-Da594 (L243, 562617), and anti-ASC-PE (HASC-71, 653904) were purchased from Biolegend (San Diego, CA, USA). Anti-NLRP3-AF700 (768319, IC7578N) was from Bio-Techne (Minneapolis, MI, USA) and, anti-NLRP1-AF647 (B-2 sc-166368) was from Santa Cruz Biotechnology (Santa Cruz, CA, USA).

### Cross-linking reagents

DMTMM (Sigma-Aldrich) is commercially available. For all cross-linking experiments, DMTMM (Sigma-Aldrich) was prepared at a 120 mg/mL concentration in 1× PBS pH 7 and stored at −80 °C as a stock solution.

### Cross-linking mass spectrometry

Immunoprecipitated NLRP1 samples were resuspended in 1xPBS, 1 mM DTT pH 7.4 buffer. All experiments were performed as biological triplicates. All samples were incubated at 37 °C while shaking at 350 rpm for 30 min. Final concentrations of 36 mM DMTMM (Sigma-Aldrich) were added to the protein samples and incubated at 37 °C with shaking at 350 rpm for 30 min. The reactions were quenched with 100 mM ammonium bicarbonate and incubated at 37 °C for 30 min. Samples were lyophilized and resuspended in 8M urea. Samples were reduced with 2.5mM TCEP incubated at 37 °C for 30 min, followed by alkylation with 5 mM iodoacetimide for 30 min in the dark. Samples were diluted to 1M urea using a stock of 50 mM ammonium bicarbonate. Trypsin Gold, MS Grade (Promega) was added at a 1:50 enzyme-to-substrate ratio and incubated overnight at 37 °C while shaking at 600 rpm. 2% (v/v) formic acid was added to acidify the samples following overnight digestion. All samples were run on reverse-phase Sep-Pak tC18 cartridges (Waters). Columns were washed with 500 μL 100% acetonitrile, washed with 1 mL 90% acetonitrile, 0.1% formic acid, sample loaded, washed again with 1mL 90% acetonitrile, 0.1% formic acid, and eluted in 50% acetonitrile, 0.1% formic acid. Ten microliters of the purified peptide fractions were injected for liquid Chromatography with tandem mass spectrometry analysis on an Eksigent 1D-NanoLC-Ultra HPLC system coupled to a Thermo Orbitrap Fusion Tribrid System. The MS was operated in data-dependent mode by selecting the five most abundant precursor ions (m/z 350–1600, charge state 3+ and above) from a preview scan and subjecting them to collision-induced dissociation (normalized collision energy = 35%, 30 ms activation). Fragment ions were detected at low resolution in the linear ion trap. Dynamic exclusion was enabled (repeat count 1, exclusion duration 30 s).

### Analysis of Mass Spectrophotometry results

All MS experiments were carried out on an Orbitrap Fusion Lumos Tribrid instrument available through the UTSW proteomics core facility. Each Thermo.raw file was converted to .mzXML format for analysis using an in-house installation of xQuest v2.1.3 (27–29). Score thresholds were set through xProphet (34–36), which uses a target/decoy model. The search parameters were set as follows: maximum number of missed cleavages = 2, peptide length = 5–50 residues, fixed modifications carbamidomethyl-Cys (mass shift = 57.02146 Da), mass shift of cross-linker = −18.010595 Da, no monolink mass specified, MS1 tolerance = 15 ppm., and MS2 tolerance = 0.2 Da for common ions and 0.3 Da for cross-link ions; search in enumeration mode. Domain-domain contact frequency plotted in GraphPad Prism v9.4.1 (Treestar).

The reduced accessible interaction space of distance-restrained NLRP1-NLRP3 complexes was computed using DisVis Webserver (30–33). For the calculations, NLRP1 and NLRP3 full-length revised structures were fetched from the AlphaFold protein structure database (accessions AF-Q9C000-F1 and AF-Q96P20-F1, respectively) (34, 35). NLRP1 was used as the reference structure and NLRP3 as the scanning protein, and using the reverse configuration lead to an impractical computational cost due to the large number of atoms in NLRP1. Distances of 0-25 Å between the Cα atoms of the crosslinked residues of the N-terminal domains of NLRP1 (residues 1-1213) and NLRP3 were included as restrictions. Sampling parameters were set to 1 Å for the grid spacing for the translational search, 9.72° for the rotational sampling interval, 3 Å threshold for the interaction radius, 200 Å^3^ for the maximum clash volume and 300 Å^3^ for the minimum interaction volume. ChimeraX version 1.7 (36) was used for the visualization.

## Statistics

Non parametric student t-test (Mann-Whitney) was used to compare data between two groups. Statistical comparisons of Flow cytometry data were performed using the Wilcoxon matched-pairs signed-rank test. Statistical analyses were performed using Prism software version 5.0a (GraphPad, San Diego, CA). All results are expressed as mean ± SD of three independent experiments and a p-value<0.05 was considered as statistically significant. Level of significance is denoted by asterisks *P < 0.05, **P < 0.01, ***P < 0.001.

## Author contributions

Conceptualization: M.D.C. Methodology: J.M.S.R., I.M.Z., A.A.G., R.A., S.B., A.G.-C., J.T.A., I.D.M., L.A.J., J.O., A.S., and M.D.C. Clinical selection and samples isolation from pateints: R.V.M., AND D.G.C. Writing (original draft): M.D.C. and A.S. Writing (review and editing): I.D.M., L.A.J., J.O., A.S. and M.D.C.

## Acknowledgments

This study was supported by the PI21/01656 grant from the Instituto de Salud Carlos III, Spain, EMC21_00033 EMERGIA program from the Junta de Andalucia, the SATOT fellowship and CNS2024-154939 from the Ministerio de Ciencia, Innovación y Universidades, Agencia Estatal de Investigación, Spain. It was also funded by the PID2021-126663NB-I0 research grant from the Spanish Ministry of Science, Innovation and Universities. JMSR is the recipient of a Postdoctoral Fellowship (RH-0085-2021) from the Consejería de Sanidad, Junta de Andalucía, Spain. This publication is part of the grant POSTD.O.C_21_00395, financed by the Junta de Andalucía /CUII and by the ESF+” (for A.G.-C.). AAG holds a Juan de la Cierva Fellowship (FJC2021-047304-I) from The Ministry of Education, Science and Sport. JTA was awarded a PhD fellowship from the Spanish Ministry of Education, Professional Training and Sport (FPU22/01094). A.S. is Wellcome Trust Senior Research Fellow. JO is supported by the grants PID2019-109276RA-I00 and PID2022-142382OB-I00 funded by MCIN/AEI/10.13039/501100011033 and by FEDER A way to make Europe, and by donations from the Spanish and Mexican Ondine Associations. J. O. was a recipient of a Leonardo Grant from the Spanish BBVA Foundation (BBM_TRA_0203) and a Ramón y Cajal Fellow of the Spanish AEI-Ministry of Science and Innovation (RYC2018-026042-I funded by MCIN/AEI/10.13039/501100011033 and by “ESF Investing in your future”). IDM’s work was supported by Grant PID2024-157414NB-I00 funded by MCIN/AEI/10.13039/501100011033 and by “ERDF A way of making Europe”, Ramon Areceś Foundation (LEUCYTO-FRA2025) and Andalusian Government (BIO-198).

## Competing interests

All other authors declare that they have no competing financial interests.

## Data and materials availability

All data needed to evaluate the conclusions in the paper are present in the paper or the Supplementary Materials.

